# Adaptive mossy cell circuit plasticity after status epilepticus

**DOI:** 10.1101/2021.04.12.439511

**Authors:** Corwin R. Butler, Gary L. Westbrook, Eric Schnell

## Abstract

Hilar mossy cells control network function in the hippocampus through both direct excitation and di-synaptic inhibition of dentate granule cells (DGCs). Substantial mossy cell loss occurs after epileptic seizures; however the contribution of surviving mossy cells to network activity in the reorganized dentate gyrus is unknown. To examine functional circuit changes after pilocarpine-induced status epilepticus, we optogenetically stimulated mossy cells in acute hippocampal slices. In control mice, activation of mossy cells produced monosynaptic excitatory and di-synaptic GABAergic currents in DGCs. In pilocarpine-treated mice, mossy cell density and excitation of DGCs were reduced in parallel, with only a minimal reduction in feedforward inhibition, enhancing the inhibition:excitation ratio. Surprisingly, mossy cell-driven excitation of parvalbumin-positive basket cells, the primary mediators of feed-forward inhibition, was maintained, indicating increased connectivity between surviving mossy cells and these targets. Our results suggest that mossy cell outputs reorganize following seizures, increasing their net inhibitory effect in the hippocampus.

## Introduction

Structural and functional alterations of hippocampal circuits in temporal lobe epilepsy manifest as imbalanced neuronal excitation and inhibition. Several mechanisms likely contribute to this imbalance, including seizure-induced cell loss, circuit reorganization such as axon sprouting, and altered receptor expression (Brooks-Kayal et al., 1998; De Lanerolle et al., 1989; Halabisky et al., 2010; Shibley and Smith, 2002; Sloviter, 1987). However, it remains unclear whether the structural alterations that occur in hilar mossy cells contribute to hyperexcitability and thus disease progression, or involve adaptive modifications that serve to maintain circuit homeostasis.

Glutamatergic hilar mossy cells give rise to bilateral long-range projections that drive direct excitation of dentate granule cells (DGCs) as well as feedforward inhibition of DGCs through local GABAergic interneurons (Hsu et al., 2016; Scharfman, 1995). Mossy cell projections are widely distributed in the hippocampus, suggesting that these cells broadly coordinate information transfer (Scharfman and Myers, 2013), and contribute to the essential role of the hippocampus in learning (Bui et al., 2018; Jinde et al., 2012). Mossy cells receive most of their excitatory and inhibitory input from neurons within the dentate gyrus and the CA3 pyramidal layer (Larimer and Strowbridge, 2008; Sun et al., 2017; Williams et al., 2007). The synaptic targets of mossy cells primarily occur in the dentate inner molecular layer (Buckmaster et al., 1996; Scharfman, 2016), albeit with projection differences between the dorsal and ventral hippocampus (Botterill et al., 2021; Houser et al., 2020).

As mossy cells are highly vulnerable to injury (Lowenstein et al., 1992; Zhang et al., 2015), alterations in mossy cell circuits after brain insults could profoundly alter hippocampal function (Santhakumar et al., 2000; Scharfman and Myers, 2013; Sloviter et al., 2003). For example, in the pilocarpine model of temporal lobe epilepsy (TLE), a large fraction of mossy cells are lost, while surviving mossy cells enlarge and receive increased excitatory input indicative of circuit reorganization (Zhang et al., 2015). The activation of residual mossy cells could either enhance or prevent seizures, depending on their net outputs, leading to different proposals regarding the residual circuitry (Botterill et al., 2019; Santhakumar et al., 2000; Sloviter, 1991; Sloviter et al., 2003). Given these opposing possibilities, a functional understanding of mossy cell output connectivity after brain injury could elucidate their contribution to hippocampal function and excitability. However, selective activation of mossy cell axons has been challenging, as electrical stimulation of these fibers in the inner molecular layer (IML) of the dentate gyrus can also activate semilunar granule cells, submammilary projections, and perforant path projections (Larimer and Strowbridge, 2008; Scharfman, 2016; Williams et al., 2007), as well as the extensive granule cell axon collaterals that sprout into the IML in epilepsy (Mello et al., 1993; Shibley and Smith, 2002; Winokur et al., 2004).

We combined genetic, viral and anatomic methods to selectively target mossy cells with channelrhodopsin *in vivo*, to facilitate selective analysis of the mossy cell circuit during an early phase of epileptogenesis. We recorded mossy cell-evoked responses from granule cells and basket cells in acute slices from healthy mice and mice following pilocarpine-induced status epilepticus. Our results suggest that the net effect of the altered mossy cell circuit following status epilepticus is inhibitory.

## Results

### Reduced monosynaptic excitation of DCGs by mossy cells after status epilepticus

We combined *in vivo* injections of channelrhodopsin-expressing viral vectors with acute brain slice electrophysiology to analyze alterations in functional mossy cell-DGC connectivity during an early phase of epileptogenesis, 3-4 weeks after pilocarpine-induced status epilepticus (SE). In this model, spontaneous seizures typically begin 2-3 weeks after SE (Mello et al., 1993; Shibley and Smith, 2002), followed by granule cell axon (mossy fiber) sprouting about 1 month after SE (Borges et al., 2003; Mello et al., 1993). We chose this time point to minimize polysynaptic activation of DGCs by sprouted mossy fibers (Hendricks et al., 2019), and only obtained single EPSCs with sharp rise times, short latency, and low jitter, suggesting that recurrent activity was not contributing to our responses.

Calcitonin Receptor-Like Receptor (Crlr)-Cre mice express Cre recombinase in hilar mossy cells of the hippocampus, with some expression in proximal CA3 cells (Jinde et al., 2012). When crossed with Rosa26-tdTomato reporter mice (Ai14), our preliminary analysis revealed significant off-target Cre-mediated recombination in pyramidal cells throughout CA3, as well as in Purkinje cells of the cerebellum. Thus, we used direct hilar injections of Cre-dependent viruses to selectively express ChR2 in mossy cells. In control mice, there was widespread ipsilateral ChR2-YFP expression in hilar mossy cells (72.8±4.4% of calretinin-positive cells co-labeled with GFP; N=4 control mice), with a dense band of bilateral IML ChR2-YFP expression corresponding to mossy cell axonal projections (Figure 1A,B). ChR2-YFP expression was not observed in GAD+ hilar interneurons or DGCs (data not shown).

**Figure 1:**
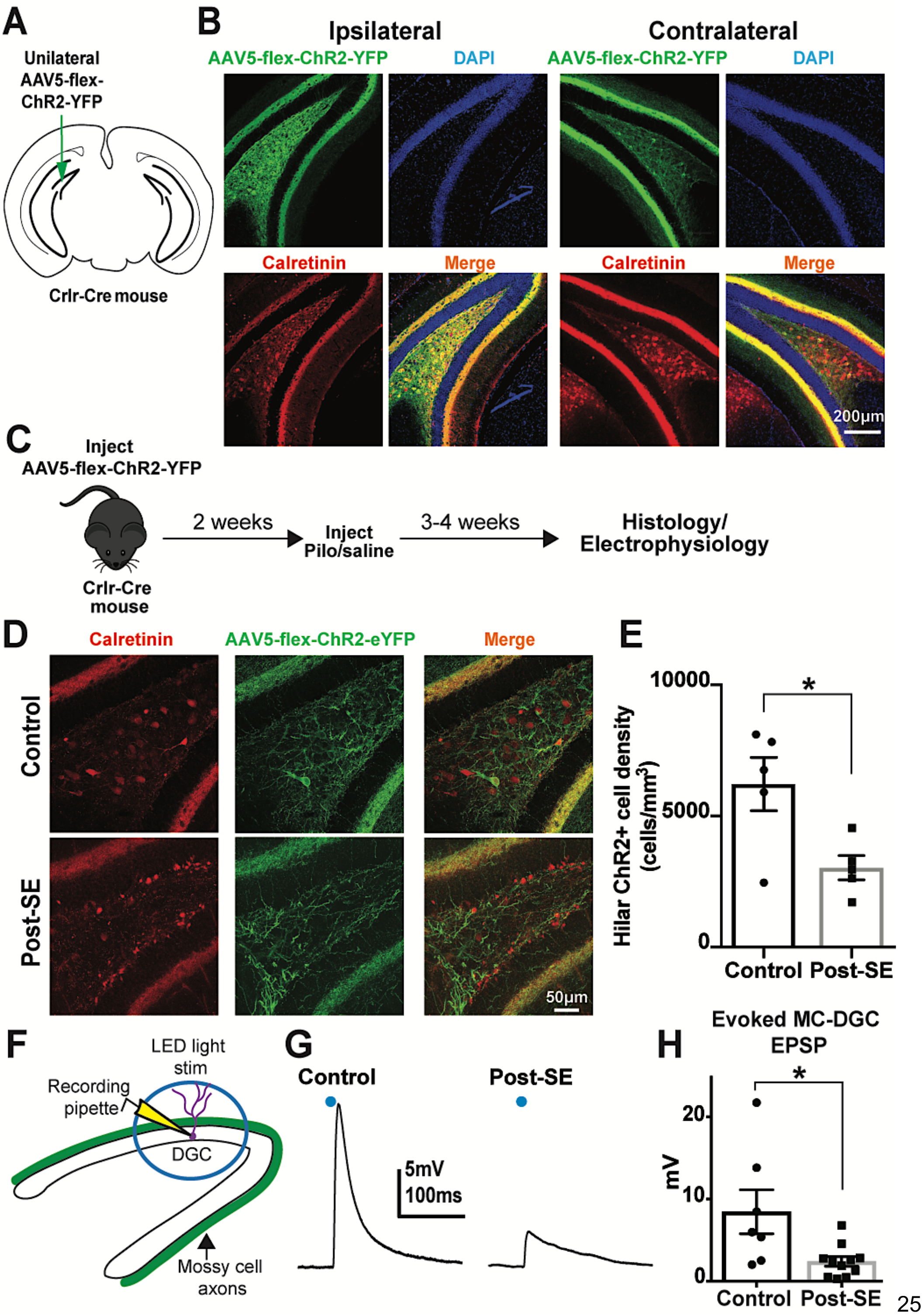
Mossy cell loss after status epilepticus reduces their excitation of dentate granule cells. **A**. Schematic of experimental design involving unilateral injection of AAV5-flex-ChR2-YFP virus into the dentate hilus of Crlr-Cre mice. **B**. Representative image of ipsilateral and contralateral ChR2-YFP expression (green), which co-labels with calretinin expressing mossy cells (red) in the ipsilateral hilus, and forms dense ipsilateral and contralateral inner molecular layer projections, consistent with mossy cell axon innervation. Nuclear stain (DAPI, blue) also shown. **C**. Experimental timeline, in which virus is injected two weeks prior to pilocarpine-induced status epilepticus (SE) or sham injections. **D**. Representative images of ChR2-YFP and calretinin expression control and post-SE mice, 3 weeks after sham/pilocarpine injections. Mossy cells are identified by their hilar location and size (arrowheads), as distinct from small calretinin-positive adult born neurons (arrows), which localize primarily to the subgranular zone and granule cell layer, and are increased after SE (lower images). **E**. Hilar ChR2+ cell density (* = p <0.05, unpaired *t* test). **F**. Schematic illustrating dentate granule cell (DGC) recording configuration during acute slice experiments and optogenetic activation of mossy cell axons. **G**. Representative recordings of optogenetically-evoked excitatory post-synaptic potentials (EPSPs) in DGCs after control or SE. **H**. Mossy cell-evoked EPSP amplitudes obtained in DGCs from control and post-SE mice (*= p < 0.05, Mann-Whitney).

ChR2+ labeled mossy cell density was reduced following pilocarpine-induced SE (control=6220±1020 cells/mm^3^, N=5 mice; post-SE=3030±460, N=5 mice: unpaired *t* test, t_8_= 2.853, p=0.0214; Figure 1D,E), consistent with hilar mossy cell loss (Zhang et al., 2015). Mossy cell loss was not due to viral injections, as calretinin labeling was similarly reduced after pilocarpine-induced SE in non-injected animals (control=3590 ±980 cells/mm^3^, N=5 mice; post-SE=1040 ±340, N=5 mice: unpaired *t* test, t_8_= 2.568, p=0.0332).

To examine direct excitatory activation of DGCs by mossy cells, we measured the excitatory postsynaptic potentials (EPSPs) evoked by optogenetic stimulation of ChR2-expressing mossy cell axons in whole-cell current clamp in the presence of the GABA_A_R antagonist SR95531 (10µM). Optogenetic stimulation reliably triggered EPSPs in control DGCs, but EPSPs were markedly reduced in DGCs from post-SE mice (control= 8.47± 2.68mV, n=7 cells (5 mice), post-SE= 2.40± 0.58mV, n=11 cells (4 mice); Mann-Whitney, p=0.02; Figure 1G,H). The latency to depolarization was not affected (control= 3.49± 0.18ms, post-SE= 3.51± 0.19ms, t_16_=0.08, p=0.94). Overall, the reduction in EPSP amplitude was similar in scale to the reduced mossy cell density after SE, demonstrating that the reduction in EPSP amplitude could be attributed to the loss of functional mossy cell innervation expected from the degree of cell loss.

The reduced light-evoked EPSP was not due to diminished optogenetic activation of surviving mossy cells in epileptic mice, as surviving mossy cells in post-SE mice reliably responded to light pulses with single action potentials, similar in efficacy to ChR2+ mossy cells in control mice (Supplemental Fig 1A,B). Input resistance was also unaltered in surviving mossy cells (control= 91.7± 27.7MΩ, post-SE= 100.1± 22.4MΩ, unpaired *t* test, t_6_=0.23, p=0.83). Additionally, we recorded spontaneous EPSPs in mossy cells from acutely prepared slices. After SE, mossy cells had an increased frequency of spontaneous EPSPs (sEPSPs) (Supplemental Fig 1C,D), similar to previous reports (Zhang et al., 2015), but no change in average sEPSP amplitude (control=1.68± 0.21mV, n=4 cells (2 mice); post-SE=1.80± 0.31mV, n=4 cells (3 mice); unpaired *t* test, t_6_=0.299, p=0.78). However, the increase in sEPSPs was not associated with increased spontaneous firing of mossy cells in our slice preparation (control=0.31± 0.17Hz, post-SE=0.10± 0.05Hz; unpaired *t* test, t_6_= 1.458, p=0.20).

### Functional properties of mossy cell-DGC synapses after SE

To gain further insights into the network consequences of mossy cell loss after SE, we examined the properties of excitatory synapses from mossy cells onto DGCs using voltage-clamp recordings. As would be expected from the reduced optogenetically-evoked potentials (Figure 1F-H), optogenetically-evoked mossy cell excitatory post synaptic currents (EPSCs) were also reduced in DGCs in post-SE mice (AMPAR-mediated EPSC; control= 45.9± 9.6pA, n=21 cells (8 mice), post-SE= 19.6± 5.1pA, n=21 cells (9 mice); Mann-Whitney, p=0.0067; Figure 2B,D). NMDAR-mediated EPSCs, recorded at a holding potential of +40 mV, were similarly reduced in post-SE DGCs (control=29.7± 5.2pA, n=16 cells, post-SE=12.6± 4.9, n=13 cells; Mann-Whitney, p=0.0048; Figure 2B,E). There was no difference in the AMPAR:NMDAR ratio between groups (control= 1.37± 0.16, post-SE=1.24± 0.20; Mann-Whitney, p=0.57; Figure 2F), indicating that there were no significant changes in the receptor complement at mossy cell-DGC excitatory synapses after SE. Passive membrane properties of DGCs were also similar between groups (Input resistance: control= 398± 46MΩ, post-SE= 487± 56MΩ, t_40_=1.24, p=0.22; series resistance: control=15.1± 1.3MΩ, post-SE=17.6± 1.8MΩ, t_40_=1.17, p=0.25).

**Figure 2:**
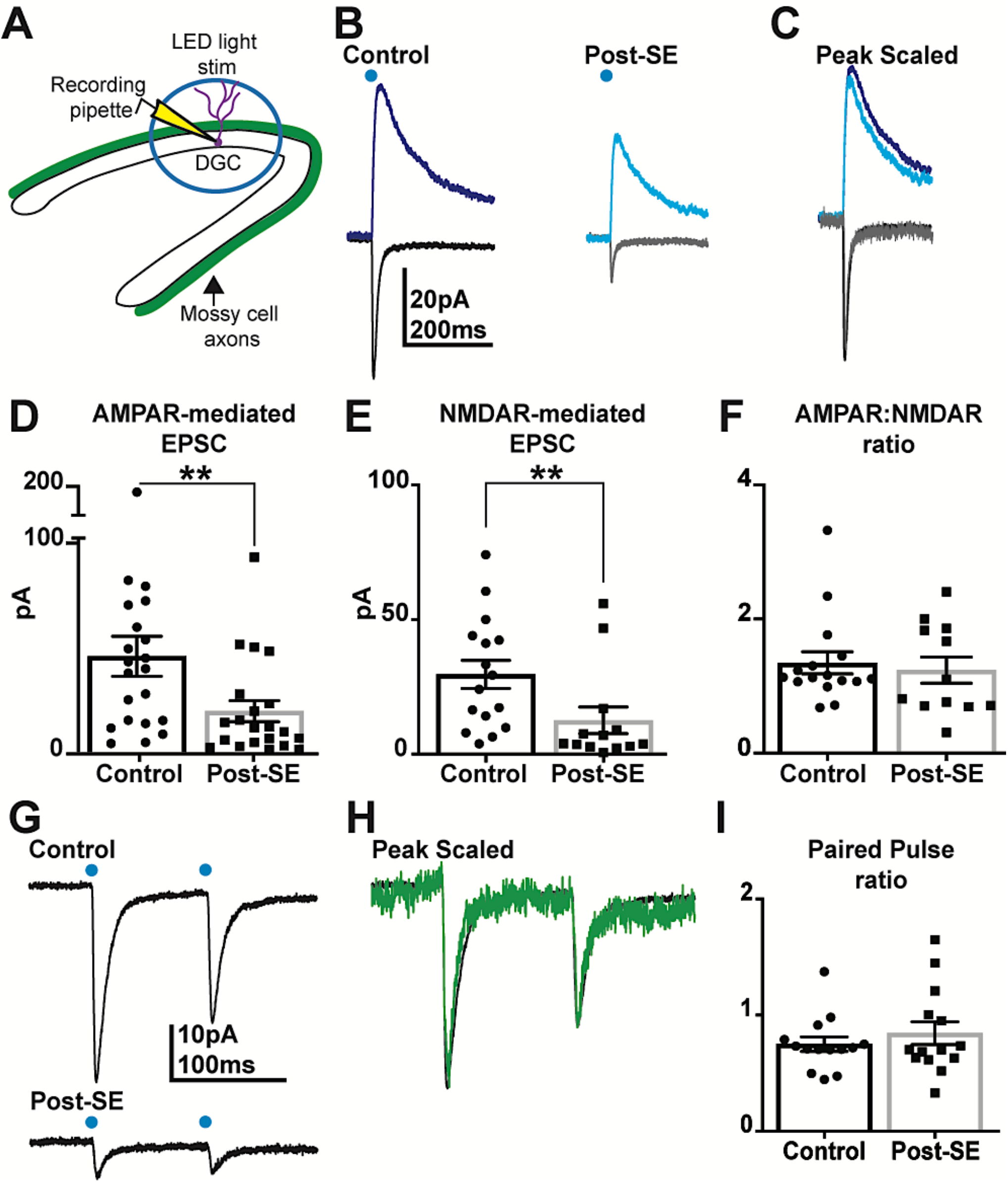
Mossy cell loss after SE does not alter receptor composition or plasticity of remaining mossy cell-DGC synapses. **A**. Schematic illustrating dentate granule cell (DGC) recording configuration during optogenetic activation of mossy cell axons. **B**. Representative excitatory post-synaptic currents in DGCs resulting from mossy cell activation in control and post-SE mice. The black trace was recorded at −70mV (AMPAR-mediated) and the blue trace was recorded at +40mV (AMPAR-+NMDAR-mediated), both with GABA_A_ receptors blocked. **C**. Peak-scaled responses (light colored traces from post-SE mice) in control and post-SE conditions. **D**. Summary data showing light-evoked AMPAR-mediated response amplitudes in DGCs from control and post-SE mice (**= p < 0.01, Mann-Whitney). **E**. Summary data showing light-evoked NMDAR-mediated response amplitudes in DGCs from control and post-SE mice (**= p < 0.01, Mann-Whitney). **F**. The ratio of AMPAR:NMDAR-mediated response amplitudes is unchanged after SE (p = 0.57, Mann-Whitney). **G**. Representative currents evoked by paired-pulse stimulation (100ms interval) in DGCs from control and post-SE mice. **H**. Peak-scaled paired-pulse responses from G (green trace from post-SE mice). **I**. Paired-pulse ratio is unchanged for mossy cell-evoked responses in DGCs from control vs post-SE mice (p = 0.95, Mann-Whitney).

Mossy cell-DGC EPSC rise and decay kinetics were also unaffected by SE (AMPA 10-90% rise-time: control=3.53± 0.35ms, post-SE= 3.25± 0.28ms, Mann-Whitney, p=0.88; AMPA decay: control=7.62± 0.39ms, post-SE=8.09± 0.63ms, Mann-Whitney, p=0.78; NMDA weighted decay: control=177± 13.9ms, post-SE= 163.2± 15.8ms, t_23_=0.64, p=0.53). Additionally, paired pulse stimulation revealed similar paired pulse depression at mossy cell-DGC synapses in post-SE mice and controls (100 ms inter-stimulus interval: control=0.75± 0.06 P2/P1 ratio, n=14 cells, post-SE=0.84± 0.10 P2/P1 ratio, n=14 cells; Mann-Whitney, p=0.95; Figure 2G-I). Together, these results suggest that reduction in population EPSC amplitudes from mossy cell-DGC synapses could be largely attributed to a decrease in the number of mossy cells, but that surviving mossy cell-DGC excitatory synapses have typical pre- and postsynaptic properties.

### Circuit plasticity shifts mossy cell-driven output toward feedforward inhibition of DGCs after SE

Di-synaptic feedforward inhibition of DGCs by mossy cells is a prominent component of the dentate mossy cell circuit (Hsu et al., 2016; Scharfman, 1995). In addition to reduced excitation of DGCs, mossy cell loss could reduce excitation of interneurons, and thus also reduce feedforward inhibition. Therefore, we optogenetically stimulated mossy cells to examine di-synaptic GABA_A_R-mediated inhibitory postsynaptic currents (IPSCs). Despite a substantial reduction in the mossy cell-driven AMPAR-mediated current after SE (57% decrease, Figure 2), there was a smaller reduction (44%) in the feedforward GABAergic input from mossy cells onto DGCs that barely reached significance (control= 41.8± 9.6pA, n=21 cells (8 mice), post-SE= 23.3± 6.4pA, n=21 cells (9 mice); Mann-Whitney, p=0.049; Figure 3B,C). When examined on a cell-by-cell basis, the ratio of inhibition to excitation (I:E ratio) driven by mossy cell activation was significantly increased after SE (GABAR/AMPAR-mediated amplitude; control=1.77± 0.58, post-SE= 3.42± 1.27; Mann-Whitney, p=0.015; Figure 3D). Thus, although both direct and indirect connectivity to DGCs was reduced after SE, there was relative preservation of the disynaptic inhibitory output from surviving mossy cells onto DGCs, shifting the net synaptic output of mossy cells towards greater inhibition of DGCs.

**Figure 3:**
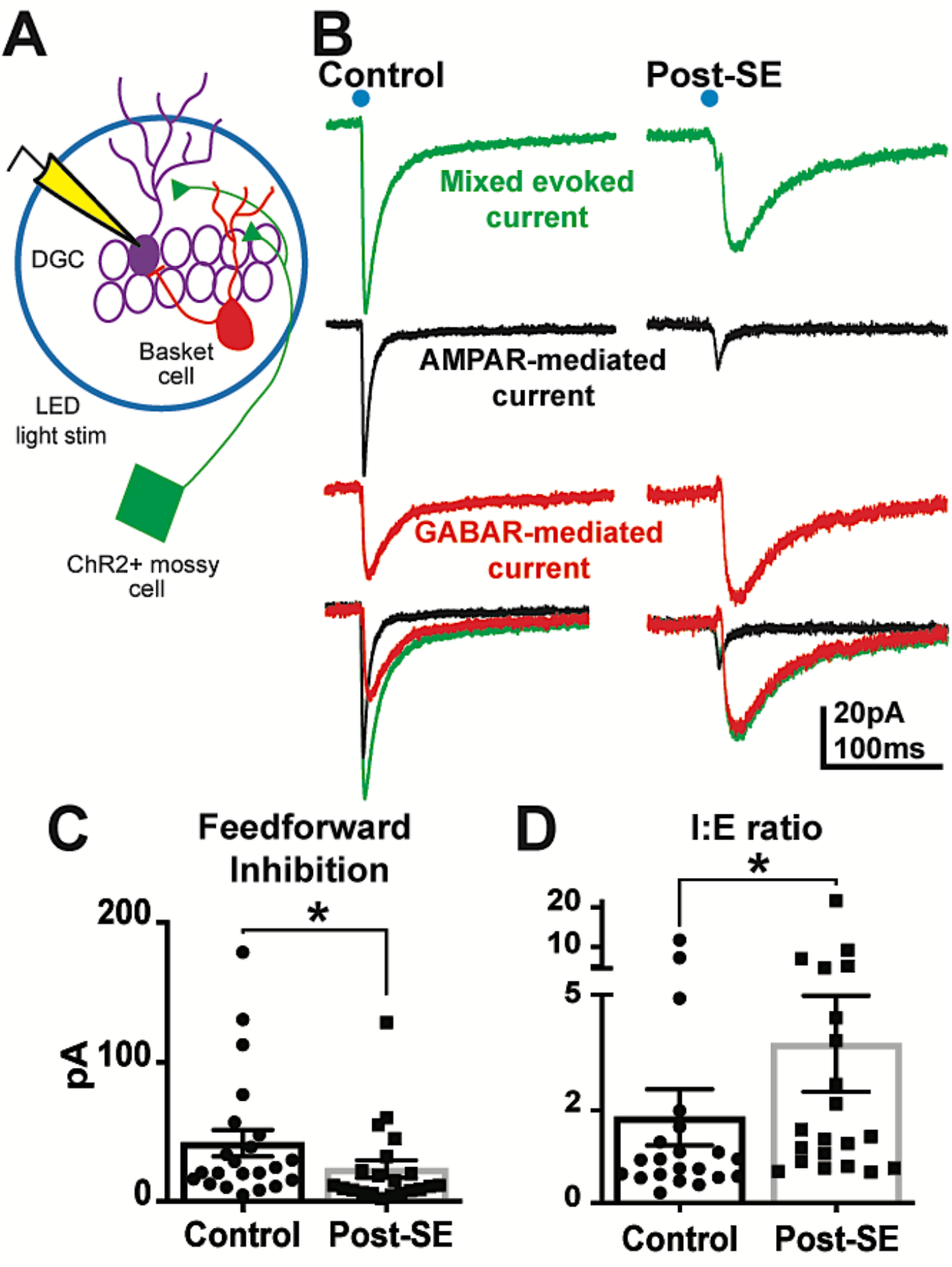
Relative preservation of mossy cell-driven feed-forward inhibition after mossy cell loss. **A**. Schematic of the dentate mossy cell microcircuit, in which mossy cells (green) directly excite DGCs (purple) and also drive di-synaptic feedforward inhibition via GABAergic interneurons, such as dentate basket cells (red). **B**. Representative optogenetically-evoked mixed currents in DGCs (−70mV, no drugs; green), as well as pharmacologically isolated, AMPAR-mediated currents (−70mV w/ 10μM SR95531; black), and GABAR-mediated (SR95531-sensitive; red) currents in DGCs from control and post-SE mice. **C**. Summary data demonstrating reduced feedforward inhibition in DGCs post-SE mice (*= p <0.05, Mann-Whitney). **D**. The ratio of inhibition:excitation in DGCs after mossy cell activation is increased after SE, indicating relative preservation of feedforward inhibition after SE (*= p <0.05, Mann-Whitney).

### Plasticity of the mossy cell to PV-basket cell circuit preserves mossy cell-mediated inhibition after SE

Parvalbumin-positive (PV+) basket cells are the primary mediators of mossy cell-mediated feedforward inhibition of DGCs (Hsu et al., 2016). These interneurons reside within the subgranular zone of the dentate gyrus, and receive excitatory innervation from mossy cells, as well as from semilunar DGCs and medial entorhinal fibers (Hsu et al., 2016; Rovira-Esteban et al., 2020). Because of the importance of PV+ basket cells in mediating feedforward inhibition of DGCs, we examined the mossy cell to PV+ basket cell circuit as a potential site of plasticity for the relative preservation of di-synaptic inhibitory mossy cell output after SE. To label PV+ basket cells for slice recording experiments, we crossed a PV-specific Cre driver line mouse (Hippenmeyer et al., 2005) with a tdTomato reporter mouse line, to identify these cells for subsequent recording.

To selectively examine the mossy cell-mediated input to PV+ basket cells, we unilaterally injected the dentate hilus of PV-Cre::tdT mice with a non-Cre-dependent channelrhodopsin virus (AAV5-Syn-Chronos-GFP), to target commissural projections of mossy cells to the contralateral dentate gyrus. This strategy allowed us to selectively stimulate mossy cell inputs while recording from genetically identified (tdTomato+) PV+ basket cells. Although these viral injections drove widespread Chronos expression in multiple cell types in the injected hemisphere, Chronos-GFP expression in the contralateral hippocampus was restricted to mossy cell axons in the IML (Fig 4A,B). Two weeks after virus injection and mossy cell labeling, mice were randomized to either control or pilocarpine-mediated SE. At the same early timepoint (3-4 weeks after SE), acute hippocampal slices were prepared to examine the connectivity of mossy cells with both DGCs and parvalbumin-tdTomato-positive (PVtdT+) basket cells using whole cell patch clamp recordings (Figure 4C).

**Figure 4:**
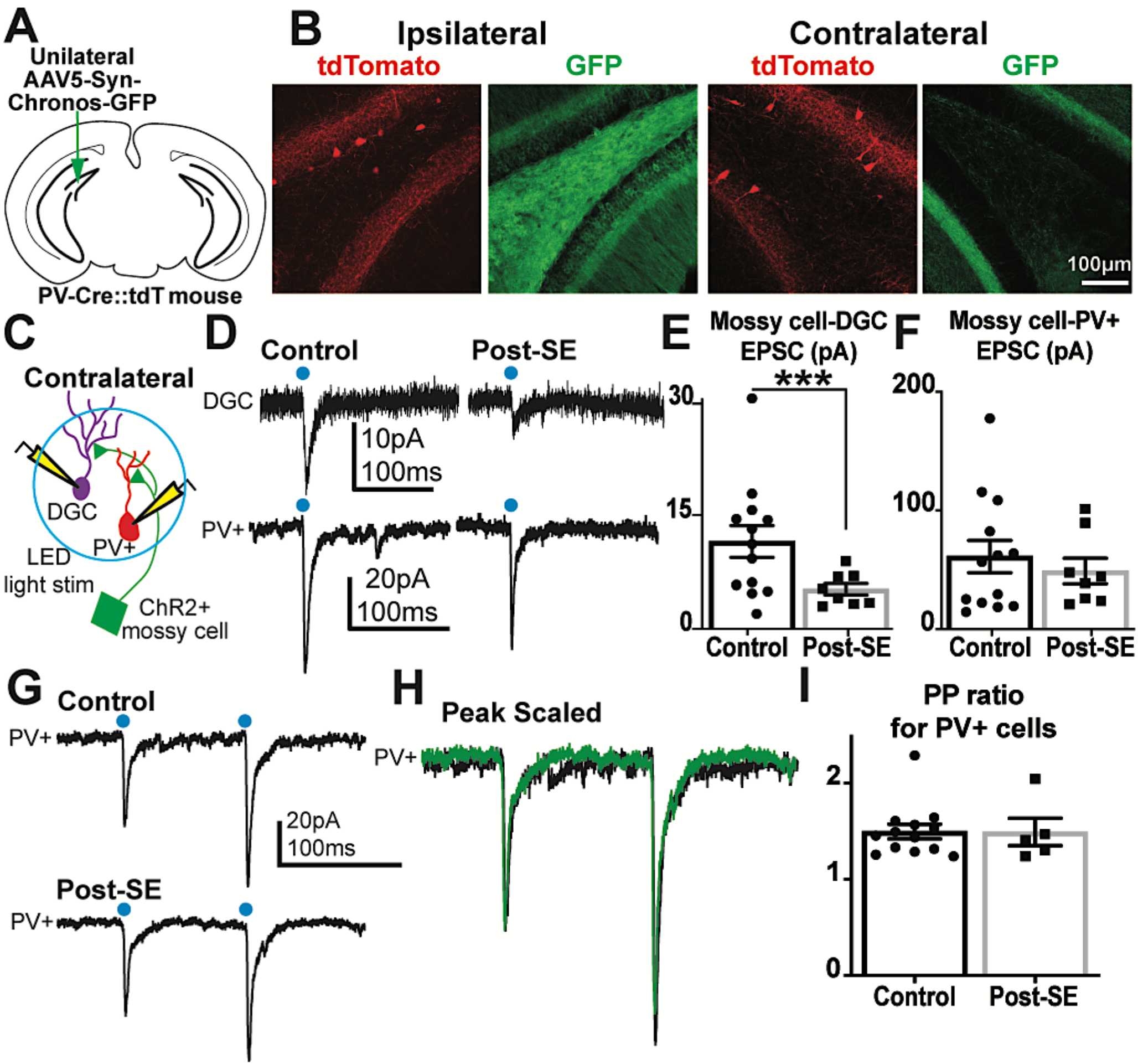
Overall mossy cell-driven excitation of PV+ basket cells is preserved despite mossy cell loss after SE. **A**. Schematic of unilateral channelrhodopsin (Chronos-GFP) virus injection in PV-Cre::tdTomato mice. **B**. Images demonstrate Chronos-GFP expression (green) in the ipsilateral dentate and the contralateral inner molecular layer, consistent with mossy cell expression. PVtdT+ basket cells shown in red. **C**. Schematic of the dual recording setup to test mossy cell connectivity onto DGCs and PVtdT+ basket cells in PV-Cre::tdTomato mice. **D**. Representative optogenetically-evoked EPSCs recorded from simultaneously-recorded PVtdT+ basket cells and DGCs in control and post-SE mice. **E**. Evoked mossy cell-DGC EPSC amplitudes in control and post-SE mice, demonstrating similarly reduced amplitudes after SE (***= p <0.001, Mann-Whitney). **F**. Evoked mossy cell-PVtdT+ EPSCs are not reduced after SE mice (p =0.92, Mann-Whitney). **G**. Representative currents evoked by paired-pulse stimulation (100ms interval) in PVtdT+ basket cells from control and post-SE mice. **H**. Peak-scaled responses from G. **I**. Paired-pulse ratio is unchanged for mossy cell-evoked responses in PVtT+ basket cells from control and post-SE mice (p = 0.70, Mann-Whitney).

Surprisingly, AMPAR-mediated synaptic input onto PVtdT+ cells was unaltered after SE (control= 61.2 ±13.6pA, n=13 cells (8 mice), post-SE=49.1 ±10.7pA, n=8 cells (6 mice); Mann-Whitney, p=0.92; Figure 4D,F). There was also no change in the paired pulse ratio (control= 1.50 ±0.08 PPR, n=13 cells (8 mice), post-SE= 1.49± 0.14 PPR, n=5 cells (4 mice); Mann-Whitney, p=0.70 Figure 4G-I), suggesting that an increase in release probability at this synapse does not compensate for the reduction in the number of mossy cells post-SE.

Consistent with prior immunohistochemistry in epileptic rodent brain (Buckmaster and Dudek, 1997; Wittner et al., 2001), we did not observe a change in PV+ basket cell density following pilocarpine-induced SE (Supplemental Figure 2 A,B). PV+ cells following SE also showed no differences in sEPSC amplitude and frequency (Supplemental Figure 2 C-E), indicating that these interneurons are present in normal number and maintain excitatory input after SE-induced reduction in the number of mossy cells.

To confirm that the maintained mossy cell-PV+ functional connectivity was not due to technical differences between the two different experimental approaches (Cre-dependent or Cre-independent vectors), or to potential variation in channelrhodopsin expression between animals, we made recordings from granule cells in the same slices as our PVtdT+ recordings (either simultaneously or sequentially), while optogenetically stimulating the same set of contralateral mossy cell projections. Consistent with prior recordings, optogenetically-evoked excitatory (AMPAR-mediated) currents in DGCs were reduced after SE (control=11.7 ± 2.1pA, n=13 cells (8 mice), post-SE= 5.3 ±0.8pA, n=8 cells (6 mice); Mann-Whitney, p=0.0004; Figure 4D,E), demonstrating a selective loss of mossy-cell mediated inputs to DGCs compared to inputs to PV+ interneurons. The functionally preserved mossy cell-PV+ basket cell circuit despite widespread mossy cell loss suggests that the overall connectivity between the remaining mossy cells and target PV+ basket cells must increase to compensate and maintain overall excitatory drive of the PV+ interneuron network, which helps explain the relative preservation of mossy cell-meditated feedforward inhibition after SE.

## Discussion

Hilar mossy cells form long-range bilateral projections in the hippocampus, and are the only intrinsic hippocampal inputs to provide widespread excitatory innervation of DGCs. Modification of mossy cell activity using chemogenetics is sufficient to alter learning and memory tasks in mice (Bui et al., 2018), highlighting the possible role of mossy cells to broadly coordinate and control hippocampal function, in particular pattern separation in the dentate gyrus and DGC phase-locking (Buckmaster and Schwartzkroin, 1994; Soltesz et al., 1993). Our results confirm that mossy cell loss in the pilocarpine model of temporal lobe epilepsy reduces overall excitatory connectivity between the mossy cell population and DGCs. Surprisingly, however, there also was a shift in the net output of the mossy cell network, with a relative preservation of mossy cell-mediated inhibitory tone in the dentate microcircuit. The maintained inhibitory tone reflects adaptive circuit plasticity during epileptogenesis, and suggests that residual mossy cell activity could have anti-seizure effects in epilepsy.

### Implications of mossy cell circuit plasticity for models of epileptogenesis

Mossy cell loss is a dominant feature of temporal lobe epilepsy in humans and in animal models (Babb et al., 1984; Blümcke et al., 2000; Lowenstein et al., 1992; Nadler et al., 1980; Obenaus et al., 1993), and has led to highly-discussed hypotheses on its contribution to hippocampal hyperexcitability.

The first set of hypotheses on the role of mossy cell loss during epileptogenesis focused on excitatory connectivity between mossy cells and DGCs. Although one might expect that mossy cell loss would decrease excitatory input to DGCs, recordings of mossy cells in other disease models suggested that mossy cell loss was coincident with increased excitatory activation of the remaining mossy cells, and contributed to elevated excitation of DGCs that was AMPAR-mediated and abrogated by isolation from hilar inputs (Santhakumar et al., 2005; Santhakumar et al., 2000; Zhang et al., 2015). Taken together, these “irritable” mossy cells were hypothesized to drive dentate hyperexcitability and potentially contribute to seizure activity (Ratzliff et al., 2002; Santhakumar et al., 2000).

The alternative “dormant basket cell hypothesis” focused explicitly on the feedforward inhibitory component of the mossy cell circuit. This hypothesis was initially based on observations in slices from electrically kindled rats, in which granule cell hyperexcitability (i.e., multiple DGC population spikes after a single perforant path stimulus) was associated with mossy cell loss and decreased granule cell inhibition (Sloviter, 1991; Sloviter et al., 2003). An inability to drive paired-pulse inhibition in slices from epileptic animals led to the suggestion that loss of excitatory input from mossy cells may have rendered the remaining GABAergic basket cells “dormant”. However, this interpretation has been controversial, as other studies have demonstrated that GABAergic hilar interneurons are also vulnerable to loss in other brain injury models (Santhakumar et al., 2000). Regardless, the dormant basket cell hypothesis suggested that mossy cell loss might contribute to epileptogenesis by reducing basket cell activation.

These two hypotheses were more recently examined in mice using two different strategies. Jinde et al. (2012) used transgenic mice to selectively express an inducible diphtheria toxin receptor in mossy cells. The specific ablation of almost all mossy cells did not cause spontaneous seizures or epilepsy when mice were followed for up to 6 weeks after ablation, although whole-cell patch clamp recordings in DGCs demonstrated a transient hyperexcitability. These observations demonstrated that isolated mossy cell loss does not cause spontaneous seizures, and that the net effect of mossy cells, even in an epileptic brain, might be to reduce hippocampal hyperexcitability. Bui et al. (2018) used *in vivo* EEG recordings of seizures in a mouse model of TLE, while optogenetically activating or inactivating the surviving mossy cell population at the beginning of potential seizure-like discharges. Optogenetic activation of surviving mossy cells actually aborted seizures, whereas silencing mossy cells increased seizure frequency in these mice. Although these data do not neatly fit into the previous hypotheses, the elevated excitability of the dentate gyrus following mossy cell ablation (Jinde et al., 2012) and net anti-seizure effect of residual mossy cells in epilepsy (Bui et al., 2018) highlight the importance of mossy cells in inhibitory control of the dentate gyrus as proposed in the dormant basket cell hypothesis.

Consistent with these *in vivo* data, our results suggest a net inhibitory effect of residual mossy cell activity. However, adaptive plasticity in the mossy cell-PV+ basket cell circuit demonstrates that basket cells are not as “dormant” as might be inferred by the degree of mossy cell loss. The net inhibitory output of the mossy cell circuit suggests that mossy cells may play an important role in maintaining inhibition of DGCs in the epileptic brain. A shift in the overall balance of this microcircuit to favor inhibition is reminiscent of reorganized inputs onto surviving inhibitory interneurons post-SE and after traumatic brain injury, which have been theorized to preserve circuit homeostasis in disease (Butler et al., 2017; Halabisky et al., 2010; Hunt et al., 2011).

The increased sEPSP frequency in surviving mossy cells after pilocarpine-induced SE fits with the notion of “irritable” mossy cells, but this enhanced activation might preserve inhibitory tone in the dentate microcircuit, as demonstrated with increased seizures when mossy cells are inhibited *in vivo* (Bui et al., 2018). Thus, perhaps mossy cells might be better described as “adaptable” rather than “irritable” in the epileptic hippocampal circuit.

### Mechanisms of mossy cell circuit plasticity

Preservation of feedforward inhibition onto DGCs despite significant mossy cell loss suggests that circuit plasticity may help maintain homeostasis after brain insults. Typically, circuit plasticity involves a combination of structural and functional adaptation. Strengthening of mossy cell-PV+ basket cell connections could be mediated through higher release probability, axon/terminal sprouting, or modification of postsynaptic receptors. The lack of change in paired pulse ratio in our experiments suggests that probability of release did not change. Thus, surviving mossy cells likely strengthen connections to target interneurons by either sprouting additional synapses or by strengthening of individual synapses. Axon sprouting by other principal neurons after brain insults contributes to increased excitatory innervation of interneurons (Butler et al., 2017; Halabisky et al., 2010; Hunt et al., 2011), and seems to be a likely candidate for this preserved mossy cell-mediated feedforward inhibition. Future work determining the mechanisms that underlie the selective plasticity for mossy cell-PV+ basket cell synapses will be important to determine how generally applicable these circuit changes are after various brain insults (i.e., traumatic brain injury and stroke), and if the enhancement of this adaptive plasticity could be a promising therapeutic strategy to increase inhibition of DGCs in epilepsy, thus reducing dentate hyperexcitability and, hopefully, seizures.

## Acknowledgements

We would like to thank members of the E.S. and G.L.W. laboratories for critical feedback and discussion. Research funding was provided by the National Institutes of Health (NIH) Grant F32-NS106732 (C.R.B), the Department of Veterans Affairs Merit Review Awards I01-BX002949 (E.S.) and I01-BX004938 (E.S.); Department of Defense Congressionally Directed Medical Research Program Award W81XWH-18-1-0598 (E.S.); NIH R21-NS102948 (Ines Koerner / E.S.) and NIH Grant P30-NS061800 (OHSU Advanced Light Microscopy Core). The contents of this manuscript do not represent the views of the US Department of Veterans Affairs or the US government.

## Author Contributions

C.R.B, G.L.W., and E.S. designed experiments; C.R.B. performed data collection and analysis; C.R.B, G.L.W., and E.S. wrote the paper.

## Inclusion and Diversity Statement

N/A

## STAR Methods

### Resource Availability

#### Lead contact

Further information and requests for resources should be directed to, and will be fulfilled by, Eric Schnell.

#### Materials availability

No new unique reagents were generated during this study.

#### Data and code availability

No new unique codes were generated during this study and any and all data will be made readily available upon request.

### Experimental Model and Subject Details

Genetically modified male mice were used in experiments, beginning at 6-8 weeks of age. We used only male mice because of the sex differences in the pilocarpine model of epilepsy, in which female mice are more resistant to pilocarpine induced status epilepticus (SE) (Buckmaster and Haney, 2012). Strains used included Calcitonin Receptor-Like Receptor (Crlr)-Cre mice (Jackson Labs #023014; Jinde et al., 2012) and Parvalbumin-IRES-Cre mice (Jackson Labs #017320; Hippenmeyer et al., 2005)) crossed with Rosa26-tdTomato marker mice (Ai14; Jackson Labs #007914; Madisen et al., 2010), and maintained as homozygous colonies. For Parvalbumin-IRES-Cre::tdTomato mice, homozygous mice from each monogenic strain were bred to produce double heterozygotic offspring. Mice were housed in the Oregon Health & Science University vivarium, with food and water provided *ad libitum* and with a 12 hr light/dark cycle. All procedures were approved by the Oregon Health & Science University Animal Care and Use Committee and followed the NIH guidelines for care and treatment of animals.

### Method Details

#### Cre-Dependent Viral Vector Delivery

To selectively label mossy cells in Crlr-Cre mice, we used unilateral intrahippocampal injections of a Cre-dependent adeno-associated virus (AAV) expressing an eYFP-tagged channelrhodopsin (pAAV-EF1a-double floxed-hChR2(H134R)-EYFP-WPRE-HGHpA, titer=3.3×10^13^ cfu/ml, 20298-AAV5, Addgene, Boston, MA), similar to a previous report (Bui et al., 2018). As our strategy to isolate mossy cell inputs onto parvalbumin-positive cells involved Cre-expressing interneurons, we isolated mossy cell inputs in these mice by infecting commissural (contralaterally-projecting) mossy cells in PV-Cre::tdTomato mice with a non-Cre-dependent channelrhodopsin virus (pAAV-Syn-Chronos-GFP, titer=5.3×10^12^ cfu/ml, 59170-AAV5, Addgene), similar to a previous report (Hsu et al., 2016). In these mice, subsequent electrophysiologic analysis was restricted to the contralateral (un-injected) hippocampus.

Intrahippocampal injections were performed as previously described (Luikart et al., 2011), with injection coordinates targeting the dentate hilus (measured from bregma: +2.35 X, −2.80 Y, and −2.20 Z). 1μL of diluted viral stock (1:3 in sterile saline) was delivered at 0.25μL/min using a calibrated Hamilton syringe (Hamilton, Franklin, MA; needle dimensions: 30g, 0.5” length, 35°angle), with the needle remaining in place for at least one minute following injection. All surgical procedures were performed under deep isoflurane anesthesia, two weeks prior to pilocarpine or vehicle injections. Mice were provided soft food with oral acetaminophen solution following surgery. Viral injections were carried out in an identical manner prior to random allocation to saline/pilocarpine injections two weeks later.

#### Induction of Status Epilepticus

We used pilocarpine to induce status epilepticus as previously described (Hendricks et al., 2017; Zhang et al., 2015). Briefly, 2 weeks following AAV injection, mice received an intraperitoneal (i.p.) injection of methyl-scopolamine (0.5mg/kg, Sigma-Aldrich, St. Louis, MO) 15-20 minutes prior to pilocarpine (325mg/kg i.p., Cayman Chemicals, Ann Arbor, MI) or saline control. Following injection, mice were visually monitored for behavioral seizures, using a modified Racine scale to score seizure severity (Racine, 1970). Mice were identified as having achieved status epilepticus after 3 seizures with a Racine score of 3 or more, followed by continuous stage 1-2 seizures. After 2 hours, seizures were terminated using diazepam (10mg/kg, i.p.). Mice were then monitored, provided soft food, and received 1mL warmed 5% dextrose solution in 0.45% saline twice daily (i.p.) until they returned to normal behavior (1-3 days). Pilocarpine-injected mice that did not reach the criteria for status epilepticus were not used for electrophysiological experiments.

Loss of hilar interneurons is common but can be highly variable in models of epilepsy, potentially due to differences in initial injury severity and subsequent development of spontaneous seizures (Jiao and Nadler, 2007). We attempted to control these variables by using a standardized time spent in status epilepticus (SE; 2hr) in pilocarpine mice and selecting a time following status (3-4 weeks post-SE) with lower spontaneous seizure likelihood, but this inherent variability could still be a factor in our data.

#### Slice preparation

3-4 weeks after pilocarpine-induced status epilepticus, mice were deeply anesthetized by isoflurane inhalation, injected i.p. with 2% 2,2,2-tribromoethanol to maintain anesthesia, and decapitated while anesthetized. The brain was removed and placed in cold (2-4°C) oxygenated NMDG-based cutting solution containing (mM): 93 NMDG, 30 NaHCO_3_, 24 glucose, 20 HEPES, 5 Na-ascorbate, 5 N-acetyl cysteine, 3 Na-pyruvate, 2.5 KCl, 2 thiourea, 1.5 NaH_2_PO_4_, 0.5 CaCl_2_, 10 MgSO_4_, and 1 kynurenic acid equilibrated with 95% O_2_-5% CO_2_ (pH 7.2-7.4). Brains were blocked, glued to a sectioning stage, and slices (300 μm) were cut in the transverse plane into cold, oxygenated NMDG-based cutting solution using a vibrating microtome (Leica VT 1200S; Leica Biosystems, Buffalo Grove, IL). Slices were transferred to a storage chamber containing oxygenated NMDG-based solution at 32-34°C for 15 minutes, then transferred to a storage chamber at room temperature with oxygenated ACSF containing (mM): 125 NaCl, 3 KCl, 2 CaCl_2_, 1.25 NaH_2_PO_4_, 25 NaHCO_3_, 1 MgCl_2_, and 25 glucose. Slices of the hippocampus from hemispheres ipsilateral and contralateral to ChR2 virus injection were used in experiments from Crlr-Cre mice, whereas only slices of the contralateral hippocampus from PV-Cre::tdT mice were used, and compared to slices from control mice that received scopolamine and saline injections.

#### Whole cell recordings

Transverse dentate gyrus hippocampal slices were transferred to a recording chamber on an upright, fixed-stage microscope equipped with infrared, differential interference contrast optics (Olympus BX50WI), and continuously perfused with ACSF. Recordings were performed at room temperature from visually identified dentate granule cells in the outer half of the granule cell layer, YFP-positive hilar mossy cells, or from tdTomato-positive basket cells near the subgranular region. Recording pipettes were pulled from borosilicate glass (TW150F; World Precision Instruments, Claremont, CA) with a P-87 puller (Sutter Instruments, Novato, CA). The intracellular solution for current clamp recordings contained (in mM): 130 KGluconate, 20 KCl, 10 HEPES, 0.1 EGTA, 4 MgATP, and 0.3 NaGTP. The intracellular solution for voltage clamp recordings contained in (mM): 113 CsGluconate, 8 NaCl, 10 EGTA, 10 HEPES, 1 MgCl_2_, 1 CaCl_2_, 3 CsOH, 0.3 MgATP and 2 NaGTP. Open tip series resistance was 3-6 MΩ. Recordings were obtained using an Axopatch 1D amplifier (Molecular Devices, Sunnyvale, CA), low-pass filtered at 10 kHz, digitized at 20 kHz with a NIDAQ (National Instruments) analog-to-digital board, and acquired using Igor Pro 5.05A (Wavemetrics) script with NIDAQmx (National Instruments) plugins. Cells were whole-cell voltage-clamped at −70mV for 5-10 min to allow equilibration of pipette and intracellular solutions prior to data collection of light evoked responses. Pulses of blue LED light (Thorlabs, 470nm, 1ms, 7.85mW/cm^2^) were delivered through a 40X water immersion objective above the YFP-positive hilar mossy cell or within the inner molecular layer for DGC or PV-tdT basket cell recordings. The GABA_A_-receptor antagonist SR95531 was used (10µM) to pharmacologically isolate excitatory synaptic input during optogenetic stimulation when indicated. The di-synaptic nature of inhibitory currents during mossy cell evoked responses was confirmed by holding DGCs at the AMPAR reversal potential (0mV) and applying the AMPAR-antagonist NBQX (10µM), which eliminated the evoked inhibitory current at 0mV as well as the total evoked response at −70mV (data not shown). N-Methyl D-Aspartate receptor (NMDAR)-mediated currents were recorded by holding DGCs at +40 mV to relieve these channels of their magnesium block, and NMDAR amplitudes were quantified 60 msec after optogenetic simulation, after the AMPAR-mediated current had ceased.

#### Immunohistochemistry and imaging

To assess viral targeting, ipsilateral hemispheres from PV-Cre::tdT mice injected with AAV5-Syn-Chronos were removed at the time of contralateral acute slice preparation and drop-fixed in 4% paraformaldehyde (PFA) in PBS for 24-48 hrs at 4°C. For mice exclusively used for immunohistochemistry, terminally anesthetized mice (2% 2,2,2-tribromoethanol, 0.8mL) were transcardially perfused with cold 0.1M PBS followed by fixative containing 4% PFA in PBS. Brains were removed and post-fixed (4% PFA in PBS) overnight at 4°C. The hippocampus was sectioned coronally (100µm) using a Leica VT 1000S vibratome and 3-4 sections (600µm interval between sections) were used for staining and imaging analysis. Sections were permeabilized with 0.4% Triton X-100 in PBS (PBS-T), blocked with 10% horse serum in PBS-T at room temperature for 1hr, and incubated in primary antibody [1:1000 goat anti-calretinin, (CG1, SWANT)] overnight at 4°C in 1.5% horse serum in PBS-T. Sections were rinsed with PBS and then incubated at room temperature for 4 hrs in PBS-T with 1:500 Alexa Fluor 488-conjugated rabbit anti-GFP (A21311, Invitrogen) and 1:500 donkey anti-goat 647 (A-21447, Invitrogen) antibodies. The tdTomato signal in PV-Cre::tdT mice was bright and visualized without antibody enhancement. Tissue was counterstained with DAPI and mounted with Fluoromount G (SouthernBiotech) onto Fisher Superfrost slides (Fisher Scientific).

We acquired images using either a Zeiss LSM 780 or LSM 900 laser scanning microscope on a motorized AxioObserver Z1 (Carl Zeiss MicroImaging). Zstack images (∼15µm) were collected with either a 5X or 20X objective. The Cell Counter plugin in FIJI (National Institutes of Health) was used to count cell densities. Imaging and quantification of cell densities were performed by an experimenter blinded to the experimental conditions.

### Quantification and Statistical Analysis

We used Igor Pro 5.05A (Wavemetrics) for curve fitting and evoked PSC analysis and Graphpad (La Jolla, CA) for additional statistical analysis. To quantify evoked responses,15-20 traces were averaged and measured for a given condition in each cell. Events characterized by a typical fast rise phase and exponential decay were selected for analysis. For all histological and electrophysiology data, normality was tested using the Shapiro-Wilk test to determine use of parametric or non-parametric statistical analysis. Data were compared using a Student’ s t test for parametric outcome measures, whereas a Mann-Whitney U test was used for non-parametric data. Data are expressed as mean ±SEM, with significance set at p<0.05. Sample sizes were selected to detect an effect size of 30-50% and a power of 0.8 with α set to p<0.05.

### Key Resources Table

| Reagent/Resource | Source | Identifier |
| --- | --- | --- |
| **Mouse Lines** |  |  |
| Calcitonin Receptor-Like Receptor (Crlr)-Cre mice | Jackson Labs | #023014 |
| Parvalbumin-IRES-Cre mice | Jackson Labs | #017320 |
| tdTomato reporter mice (Ai14) | Jackson Labs | #007914 |
| **Drugs/Electrophysiology Reagents** |  |  |
| 2% 2,2,2-tribromoethanol | Sigma | T48402 |
| SR95531 | Abcam | ab120042 |
| NBQX | Abcam | ab120046 |
| Methyl-scopolamine | Sigma | S8502 |
| Pilocarpine | Cayman Chemicals | 14487 |
| Diazepam | Hospira Inc. | NDC 0409-3213-12 |
| **Viral Vectors** |  |  |
| AAV5-EF1a-double floxed-hChR2(H134R)-EYFP | Addgene | 20298-AAV5 |
| AAV5-Syn-Chronos-GFP | Addgene | 59170-AAV5 |

| **Antibodies/Immunohistochemical Reagents** |  |  |
| --- | --- | --- |
| Goat-anti-calretinin | SWANT | CG1 |
| Alexa Fluor 488-conjugated rabbit anti-GFP | Invitrogen | A-21311 |
| Donkey anti-goat Alexa Fluor 647 | Invitrogen | A-21447 |
| DAPI | Cayman Chemicals | 14285 |
| Fluoromount-G | Southern Biotech | 0100-01 |
| SuperFrost Slides | Fisher Scientific | 22-037-246 |
| **Software** |  |  |
| Graphpad Prism | Graphpad Software Inc. | https://www.graphpad.com |
| Igor Pro 5.05 | Wavemetrics | https://www.wavemetric.com |

## Supplemental Information

**Supplemental Figure 1:**
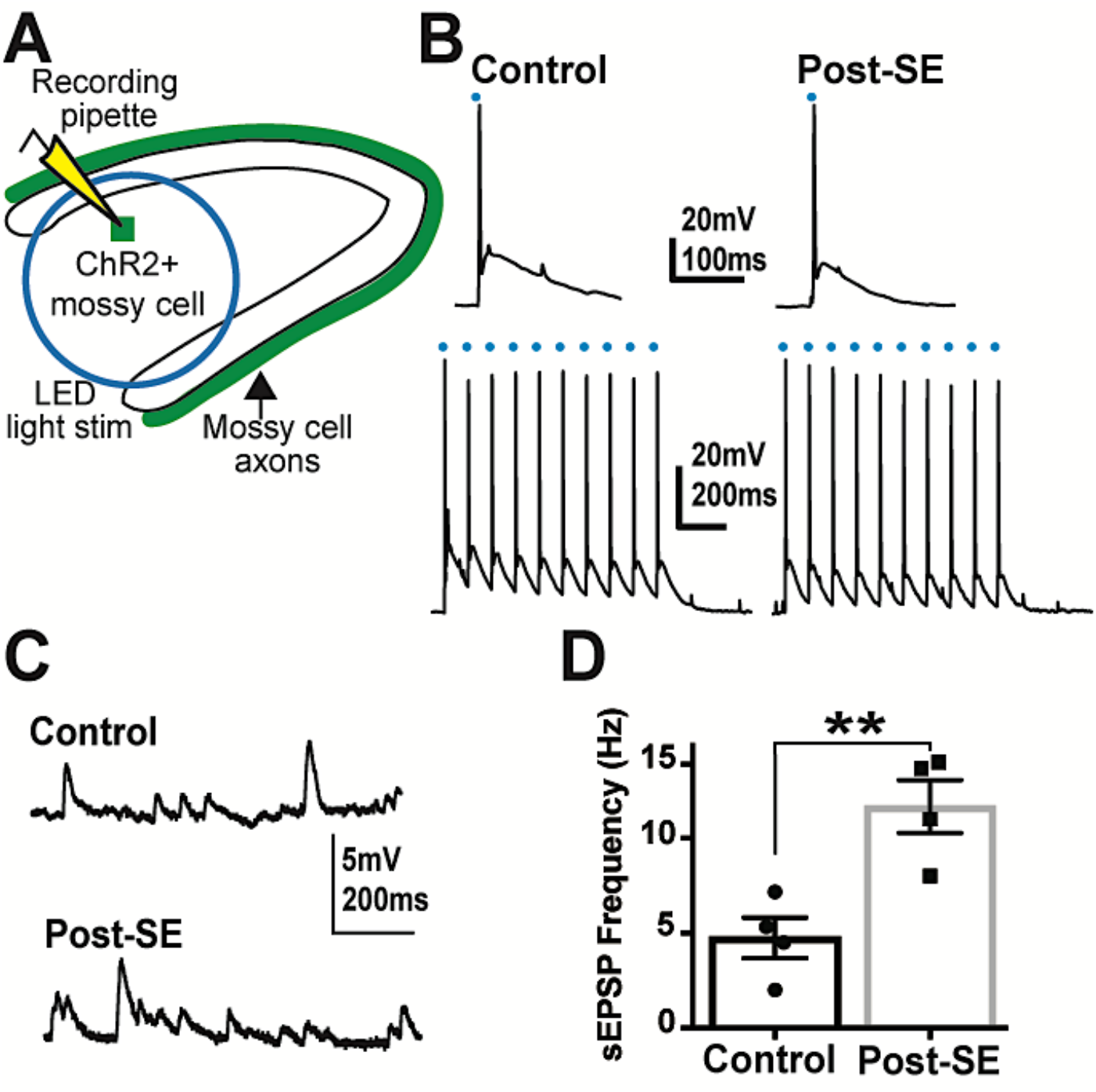
Characterization of ChR2-expressing mossy cells in control and post-SE mice. **A**. Schematic of recording setup. **B**. LED-evoked action potentials in ChR2+ mossy cells after single pulses or 10 Hz trains of 1 msec light stimulation, demonstrating similar activation of cells from control and post-SE mice. **C**. Representative spontaneous excitatory post-synaptic potential (sEPSP) recordings from ChR2+ mossy cells from control and post-SE mice in the absence of light stimulation. **D**. ChR2+ mossy cells from post-SE mice have an increased sEPSP frequency relative to controls (control= 4.75 ±1.07Hz, n=4 cells (2 mice); post-SE= 11.67 ±1.39Hz, n=4 cells (3 mice); t_6_= 3.932; p=0.0077, unpaired *t* test).

**Supplemental Figure 2.**
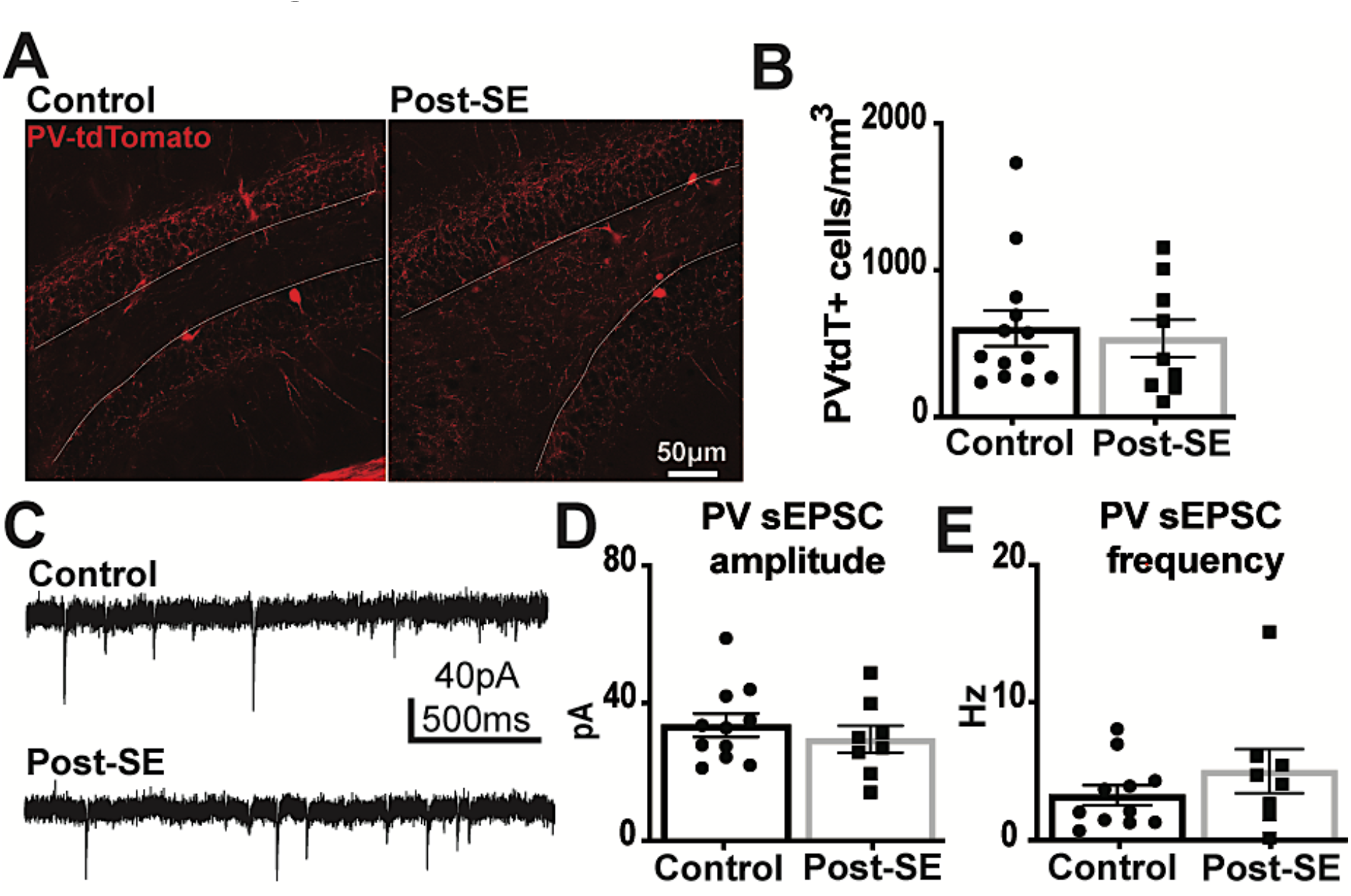
Cell density and spontaneous excitatory synaptic inputs onto PV+ basket cells are similar between control and post-SE mice. **A**. Images of PVtdT+ cells in the dentate gyrus of control and post-SE mice (white line denotes the granule cell layer / hilus border). **B**. PVtdT+ basket cell density is not different between control and post-SE mice (control= 602.9 ±121.8 PV-tdT+ cells/mm^3^, N=13 mice, post-SE= 535.1 ±127.4 PV-tdT+ cells/mm^3^, N=9 mice; p=0.60, Mann-Whitney). **C**. Representative spontaneous excitatory post-synaptic current (sEPSC) recordings in PVtdt+ cells from control and post-SE mice in the absence of light stimulation. **D**. sEPSC amplitude in PVtdT+ basket cells from control and post-SE mice is unaltered (control=33.5 ± 3.4pA, n=11 cells/7 mice, post-SE= 29.5 ±3.9pA, n=8 cells/6 mice; t_17_= 0.777; p=0.45, unpaired *t* test). **E**. sEPSC frequency is similar in PVtdt+ basket cells from control and post-SE, demonstrating that surviving PVtdT+ basket cells are not dormant in post-SE mice (control=3.23 ± 0.74Hz, n=11 cells/7 mice, post-SE= 4.99 ±1.61Hz, n=8 cells/6 mice; p=0.35, Mann-Whitney).

